## supplemental figure for "Descending locus coeruleus noradrenergic signaling to spinal astrocyte subset is required for stress-induced mechanical pain hypersensitivity"

**This PDF file includes:**
Figures S1 to S8

**Supplemental Figures**

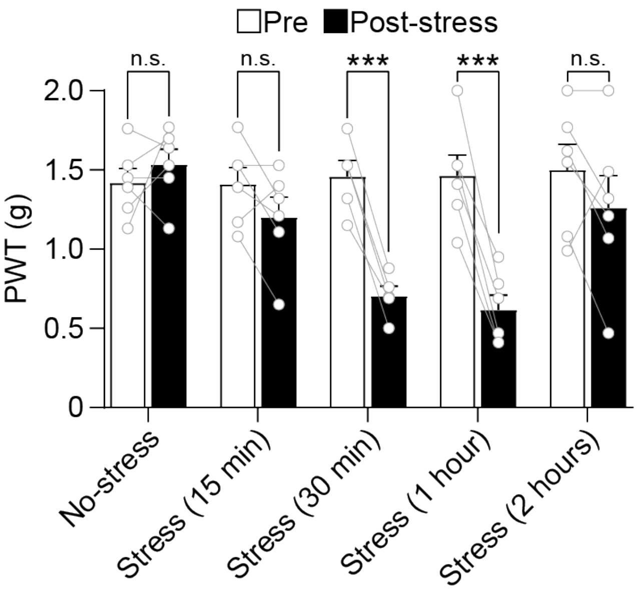


**Figure S1. Mechanical hypersensitivity is weak in mice with longer exposure (2 hours) to restraint stress.**

PWT change at 30 min after various exposure periods of restraint stress (15 min, 30 min, 1 hour, and 2 hours) in wild-type mice (*n* = 5 mice; two-tailed paired *t*-test; ***$P < 0.001$; n.s., not significant vs. pre group). Data represent mean ± SEM.

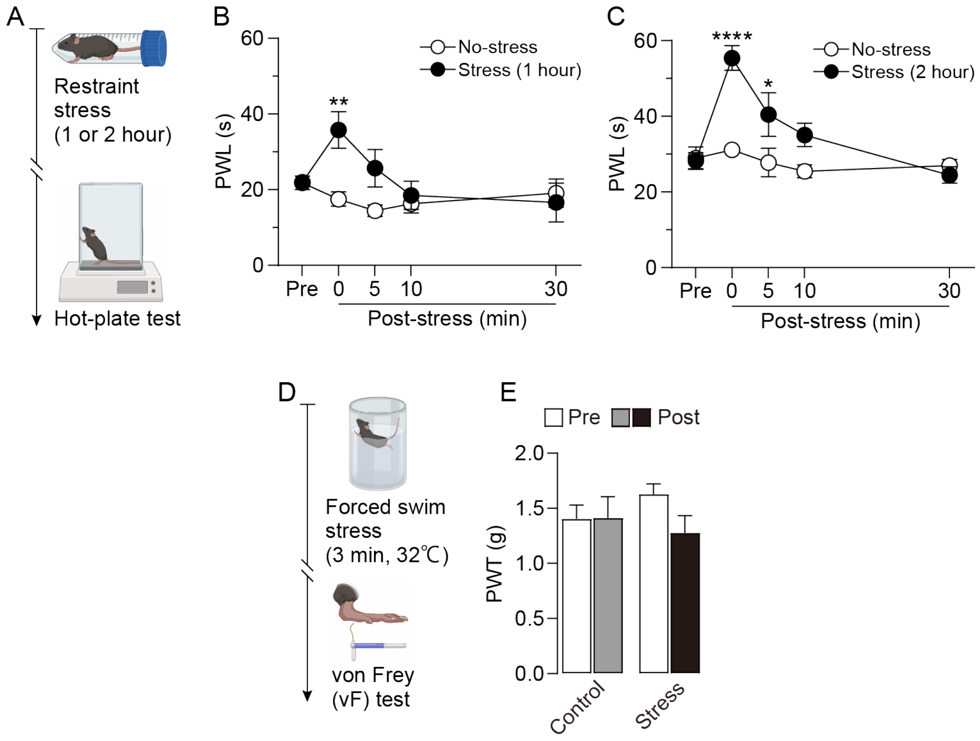


**Figure S2. Behavioral responses to thermal and mechanical stimuli in stress models.**

(A) Schematic illustration of an experiment to investigate the effects of acute exposure to restraint stress (1 and 2 hours) on thermosensory behavior in mice, using the hot-plate test. (B and C) Change in paw withdrawal latency (PWL) measured by noxious heat stimulation in wild-type mice after restraint stress for 1 hour (B) or 2 hours (C) (B: $n$ = 10 mice per group) (C: $n$ = 5 mice per group) (two-way ANOVA with post hoc Bonferroni's multiple comparisons test; *$P < 0.05$, **$P < 0.01$, ****$P < 0.0001$ vs. no-stress group). (D) Schematic illustration of an experiment to investigate the effects of acute exposure to forced swim stress (3 min, 32°C) on mechanosensory behavior in mice, assessed with the von Frey test. (E) PWT measured by mechanical stimulation in wild-type mice after forced swim stress ($n$ = 5 mice per group; two-tailed paired t-test and two-way ANOVA with post hoc Bonferroni's multiple comparisons test). Data represent mean ± SEM.

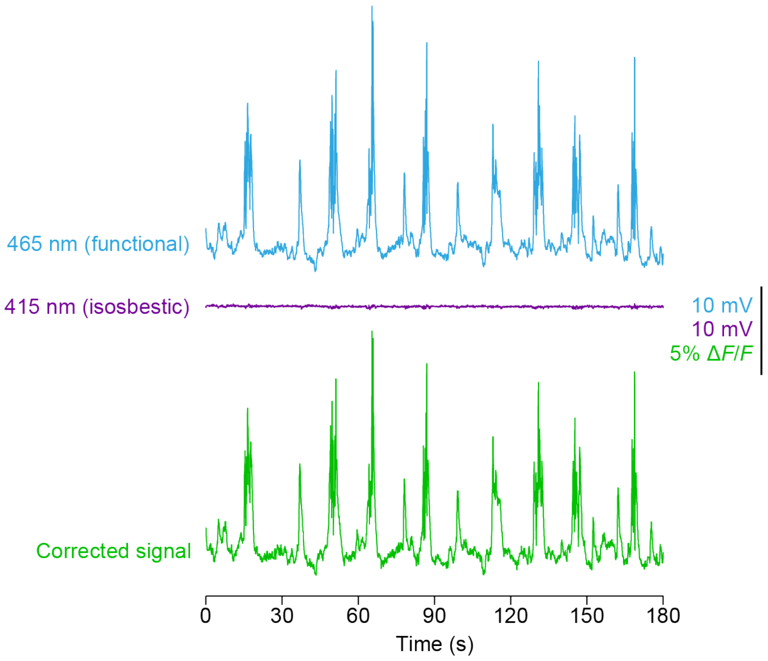


**Figure S3. Raw fluorescent signals in LC-NA neurons during restraint stress (*in vivo* fiber photometry).**

Representative traces of GCaMP6s signals in LC-NA neurons during restraint stress. Traces shown at the top (blue), middle (purple), and bottom (green) indicate 465-nm, 415-nm, and corrected fluorescent signals, respectively.

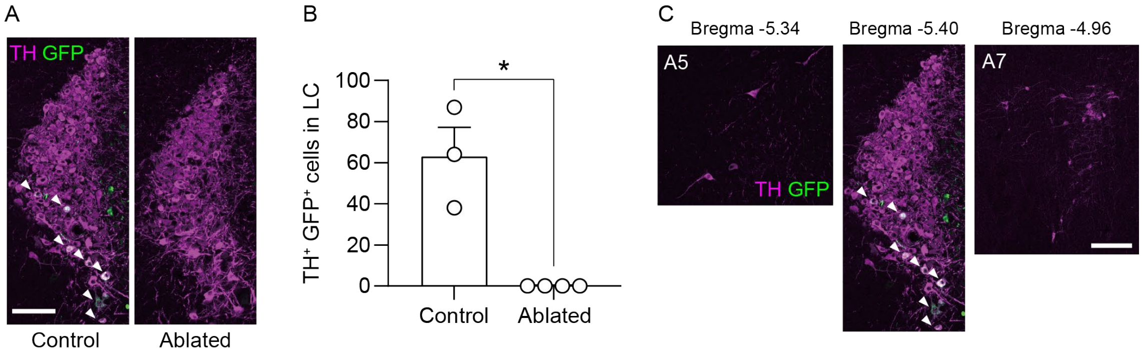


**Figure S4. Specific expression and ablation of $LC^{\rightarrow SDH}$-NA neurons.**

(A) Representative of $LC^{\rightarrow SDH}$ -NA neurons in DTR-expressing mice treated with vehicle (control) and DTX (ablated). Scale bar, 100 µm. (B) Number of $TH^{+}$ $GFP^{+}$ cells in multiple sections of the anterior and posterior regions of the LC ($n$ = 3–4 mice; Mann Whitney test; *$P < 0.05$). (C) Representative of $LC^{\rightarrow SDH}$ -NA neurons in DTR-expressing mice (a retrograde AAV vector incorporating Cre injected into the SDH and an AAV vector incorporating DTR (fused with EGFP) injected into the LC). DTR expression was not observed in the A5 or A7 regions, indicating that DTR expression was specific to the A6 (LC) region (this image is the same as Control in panel A). Scale bar, 100 µm. Data show the mean ± SEM.

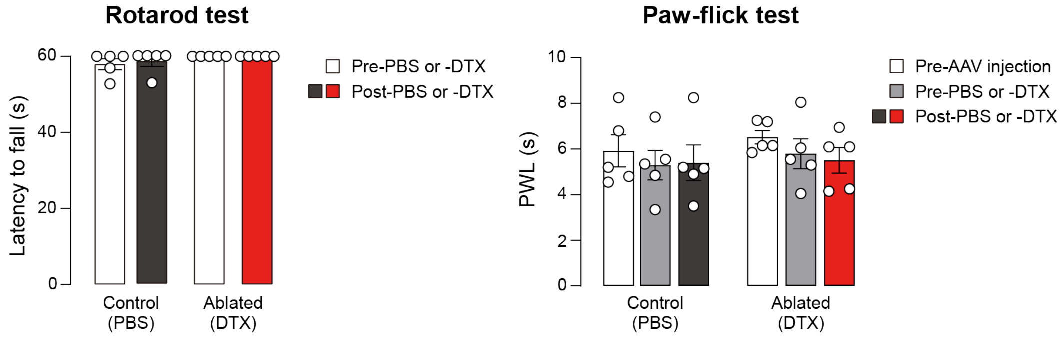


**Figure S5. Ablation of LC$^{\rightarrow SDH}$ -NA neurons does not affect motor function or thermal sensation.**

Latency to fall in the rotarod test (left) and paw withdrawal latency (PWL) in the paw-flick test (right) using mice with DTR expression in LC$^{\rightarrow SDH}$ -NA neurons before and after injection of DTX or PBS (*n* = 5 mice per group; two-tailed paired t-test and two-way repeated measures ANOVA with Bonferroni's multiple comparisons test). Data show the mean ± SEM.

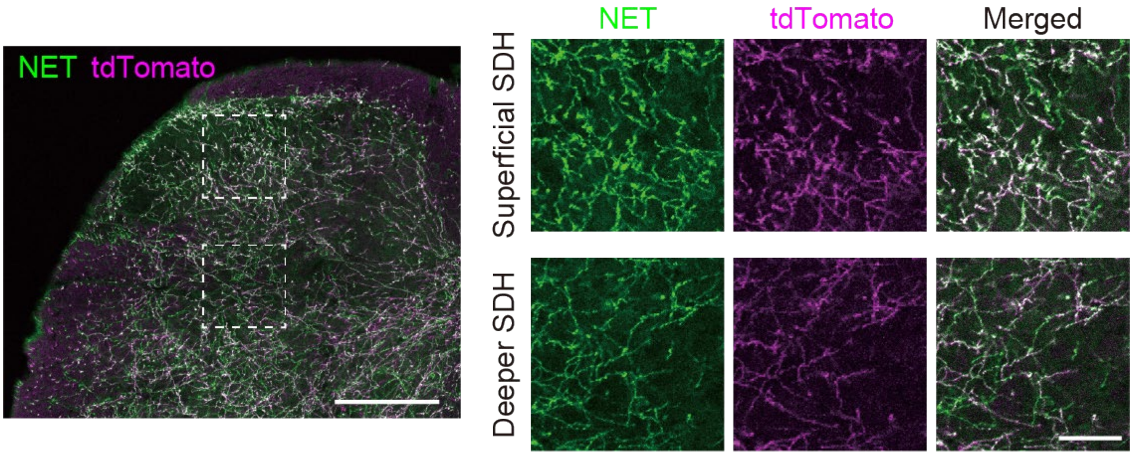


**Figure S6. Expression of ChrimsonR in the superficial and deeper SDH.** Representative images of NET (green) and tdTomato (magenta) expression in the SDH at 3 weeks after intra-LC injection of AAV-FLEx[ChrimsonR-tdTomato] in *NET-Cre* mice. Scale bar, 200 µm (left) and 50 µm (right).

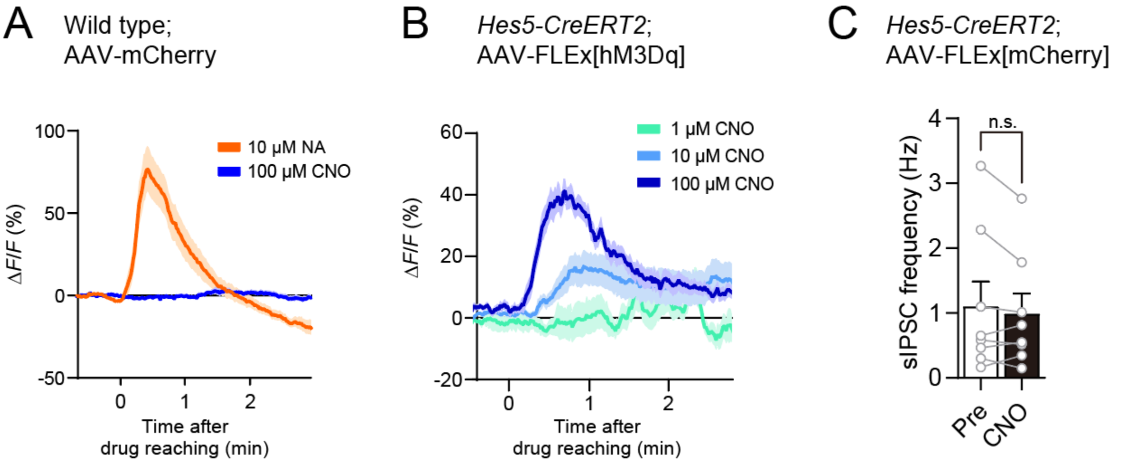


**Figure S7. Effects of CNO on $Ca^{2+}$ response in astrocytes and IPSCs in SG neurons in spinal cord slices from mice with or without hM3Dq expression.**

(A) Representative images of the astrocytic $Ca^{2+}$ response by NA (10 μM) or CNO (100 μM). Fluorescence intensity of GCaMP6m in wild-type;AAV-mCherry mice with GCaMP6m expression in SDH astrocytes. (B) Effect of CNO (1, 10, and 100 μM) on the astrocytic $Ca^{2+}$ response in *Hes5-CreERT2*;AAV-FLEx[hM3Dq] mice with GCaMP6m expression in SDH astrocytes. (C) Frequency of sIPSCs in SG neurons in the SDH from *Hes5-CreERT2*;AAV-mCherry mice treated with TAM [Pre and CNO: before and after bath application of CNO (100 μM), respectively] ($n$ = 8 cells from 8 mice; Wilcoxon signed-rank test; p = 0.5781). Data show the mean ± SEM.

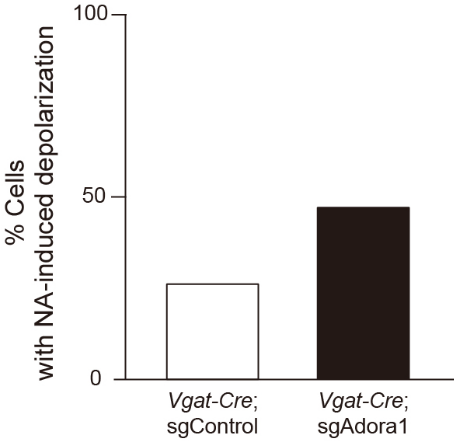


**Figure S8. Knockdown of $A_1$Rs in Vgat$^+$ neurons tends to increase the proportion of Vgat$^+$ neurons with NA-induced depolarizing responses.**

Vgat$^+$ neurons were recorded in spinal cord slices of *Vgat-Cre* mice with AAV-FLEx[SaCas9] and AAV-FLEx[mCherry]-U6-sgAdora1 or -FLEx[mCherry]-U6-sgControl, and were classified as depolarizing based on the peak change in membrane potential following bath application of NA (sgControl: *n* = 38 from 13 mice; sgAdora1: *n* = 17 from 7 mice; Fisher's exact test; p = 0.2127).
